# Caspase cleavage of Influenza A virus M2 disrupts M2-LC3 interaction and regulates virion production

**DOI:** 10.1101/2024.04.04.588074

**Authors:** Carmen Figueras-Novoa, Masato Akutsu, Daichi Murata, Ming Jiang, Beatriz Montaner, Christelle Dubois, Avinash Shenoy, Rupert Beale

## Abstract

Influenza A virus (IAV) Matrix 2 (M2) protein is an ion channel, required for efficient viral entry and egress. M2 interacts with the small ubiquitin-like LC3 protein through a cytoplasmic C-terminal LC3 interacting region (LIR). Here, we report that M2 is cleaved by caspases, abolishing the M2-LC3 interaction. A crystal structure of the M2 LIR in complex with LC3 indicates the caspase cleavage tetrapeptide motif (_82_SAVD_85_) is an unstructured linear motif that does not overlap with the LIR. Furthermore, an IAV mutant expressing a permanently truncated M2, mimicking caspase cleavage, is impaired in M2 plasma membrane transport and produces a significantly attenuated virus. Our results reveal a dynamic regulation of the M2-LC3 interaction by caspases. This highlights the role of host proteases in regulating IAV exit, relating virion production with host cell state.

## Introduction

Influenza A virus (IAV) is an important human and animal pathogen. As well as being a major pandemic threat, seasonal IAV results in an estimated 290,000 to 650,000 respiratory deaths per year [1]. IAV is an eight segment genome negative-strand RNA virus which belongs to the *Orthomyxoviridae* family [2]. The IAV envelope is derived from the host cell plasma membrane, containing viral hemagglutinin (HA) and neuraminidase (NA) proteins [3]. A small amount of viral Matrix 2 (M2) is also incorporated into virions. M2 is a single-pass membrane protein, comprising an external N-terminal domain, transmembrane domain, and a cytosolic C-terminal domain. M2 forms a tetrameric pH-regulated, proton selective ion channel, activated by low exterior pH, and is a target for antiviral drugs [4–7]. The M2 protonophore activity is required for both viral entry and egress, where premature activation of HA would otherwise occur during trafficking through the secretory pathway [8–10].

De-acidification of the secretory pathway results in erroneously neutral compartments. Erroneous pH has been proposed to induce recruitment of ATG16L1, a component of the core autophagy machinery, by V-ATPases. The E3-ubiquitin-ligase like ATG16L1 complex induces lipidation of the ubiquitin-like ATG8 molecules, comprising the LC3 and GABARAP proteins. This conjugation of ATG8s to single membranes (CASM) is distinct from canonical macroautophagy [11, 12]. CASM depends on an interaction between the WD40 domain of ATG16L1 and V-ATPases, which is strongly inducible by M2 proton channel activity [12–15].

Although CASM activation during IAV infection is mediated through M2 proton channel activity, M2 further modulates CASM by the presence of an LC3-interacting region (LIR) motif in its C-terminal domain [11]. The M2 LIR motif directly binds to ATG8s and promotes their relocalisation to the plasma membrane. When point mutations are introduced in the M2 LIR, mutant viruses exhibit attenuated stability and reduced filamentous budding. [11]. The presence of a highly conserved LIR motif in M2 suggests that IAV not only triggers CASM as a cellular response, but also utilises it during egress.

Caspases, a family of cysteine proteases, are responsible for the execution of cell death pathways such as apoptosis and pyroptosis [16]. Caspases recognise four amino acid long motifs that end in an aspartic acid, which is essential for substrate cleavage. Many caspase substrates have been identified, including viral and autophagy-related proteins [17, 18]. M2 has been proposed to be a target of caspase cleavage [19]. We speculated that the role of caspase cleavage of M2 might be to remove the LIR motif in response to cellular caspase activation. Here we experimentally identify a caspase cleavage motif in the C-terminal domain of M2. Cleavage removes the LIR and disrupts the M2-LC3 interaction. Permanently truncated IAV M2 mutant virus was substantially attenuated, highlighting the importance of the M2-LC3 interaction and its modulation through cleavage.

## Results

### M2 is cleaved at _82_SAVD_85_ motif by caspases

To study M2 cleavage during infection, PMA-differentiated THP-1 macrophage-like cells were infected with IAV, and samples collected in two-hour intervals between 8 and 16 hours (Figure 1A). We observed the appearance of a faster migrating M2 which we hypothesised could correspond to an M2 cleavage product. We noted concomitant PARP-1 cleavage, indicating caspase activation and subsequent cleavage of substrates (Figure 1A). To further analyse the role of caspases in M2 cleavage, THP-1 cells were infected with IAV and treated with Z-VAD-FMK, a pan-caspase inhibitor. We observed a lower molecular weight M2 band formed at late time points (Figure 1A-B). However, the band corresponding to cleaved M2 was not present upon Z-VAD-FMK treatment, indicating that M2 is cleaved by caspases (Figure 1B).

**Figure 1.**
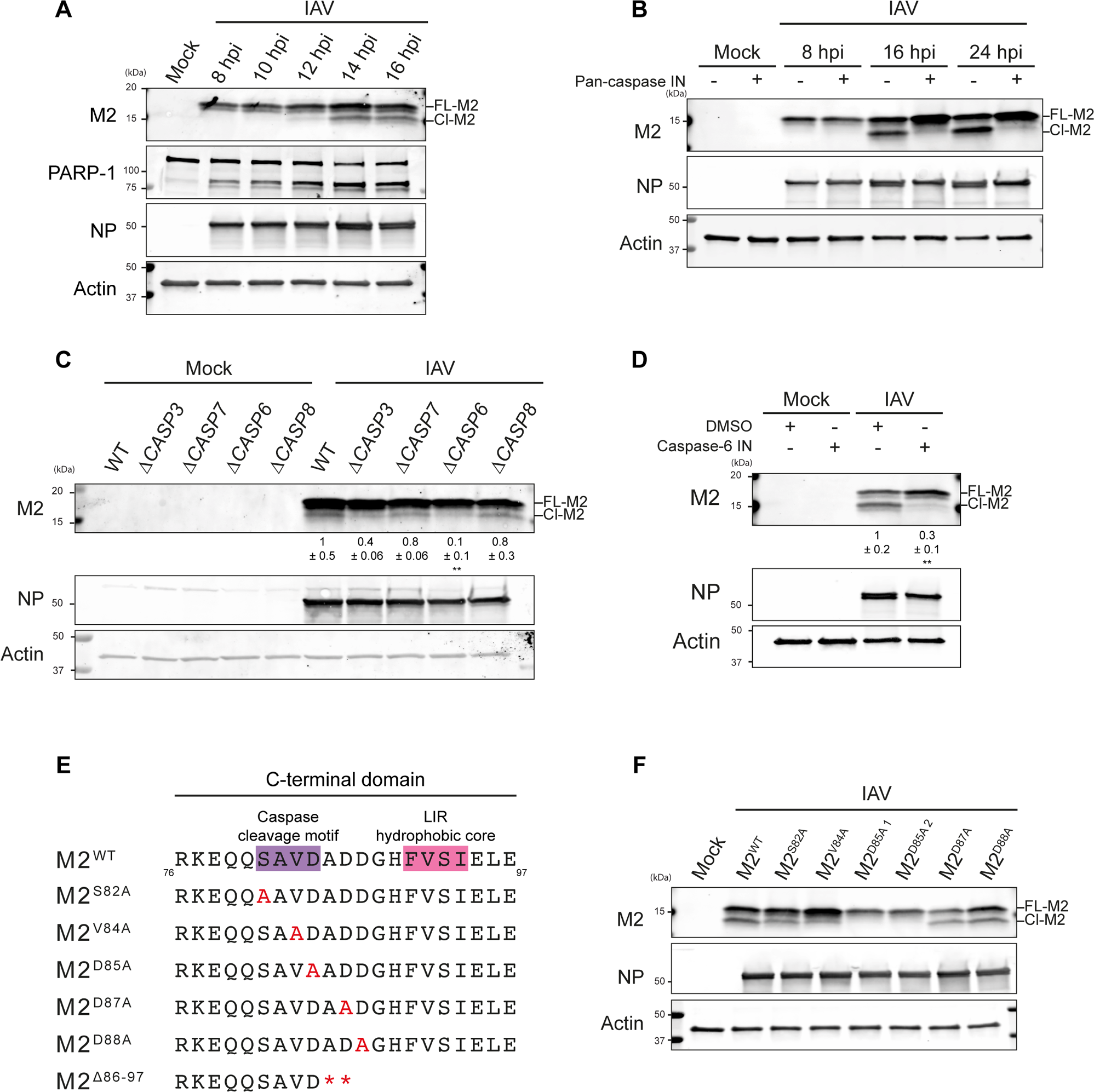
M2 is cleaved at _82_SAVD_85_ motif by caspases. **A,** Representative immunoblots of lysates of THP-1 cells infected with IAV PR8 for 8, 10, 12, 14, 16 hours with an MOI of 10. **B,** Representative immunoblots of lysates THP-1 cells infected with IAV PR8 for the indicated duration with an MOI of 10. Indicated samples were treated with DMSO as a control, or 20 µM Z-VAD-FMK as indicated. **C,** Representative immunoblots of lysates of wild type (WT), Δ*CASP3*, Δ*CASP7*, Δ*CASP6*, and Δ*CASP8* HAP1 cells. The indicated samples were from cells infected for 16 hrs with IAV PR8 at an MOI of 10. The ratio of cleaved M2 to full length M2 was quantified. Mean ± SD of n = 3 shown. **: P ≤ 0.01. Ordinary one-way ANOVA with Bartlett’s multiple comparisons. **D,** Representative immunoblots of lysates of THP-1 cells infected with IAV PR8 for 24 hours. Indicated samples were treated with DMSO as a control, or 20 µM caspase-6 inhibitor (Z-VEID-FMK) as indicated. The ratio of cleaved M2 to full length M2 was quantified. Mean ± SD of n = 3 shown. **: P ≤ 0.01. Unpaired *t* test. **E,** Diagram depicting C-terminal region of IAV PR8 (strain A/Puerto Rico/8/1934). The caspase cleavage motif is highlighted in purple and the LIR hydrophobic core in pink. Amino acid changes for mutants mentioned are highlighted in red. **F,** Representative immunoblots of lysates of THP-1 cells infected with IAV PR8 single amino acid mutants in and around the C-terminal caspase cleavage motif with an MOI of 10.

To identify caspases responsible for M2 cleavage, we used HAP1 cells deficient for the three executioner caspases (caspase-3, −7 and −6) and initiator caspase, caspase-8 [20]. We found that Δ*CASP6* cells had substantially reduced levels of cleaved M2 upon IAV infection (Figure 1C, Figure S1A). To confirm the role of caspase-6 in M2 cleavage, Δ*CASP6* cells were reconstituted to stably express caspase-6 (Δ*CASP6*+CASP6; Figure S1B). Expression of caspase-6 in Δ*CASP6* cells rescued M2 to cleavage to levels seen in the parental cell line (Figure S1B-C).

To further test the role of caspases in M2 cleavage, THP-1 cells were infected and subsequently treated with the caspase-6 inhibitor Z-VEID-FMK followed by immunoblots for M2 cleavage. Compared to vehicle-treated THP-1 cells infected with IAV, Z-VEID-FMK treatment significantly reduced the abundance of cleaved M2 (Figure 1D, Figure S1D). This supported a role for caspase-6 in M2 cleavage. Caspase-3 (Z-DEVD-FMK) and caspase-8 (Z-IETD-FMK) inhibitors were also used to assess the roles of these caspases in M2 cleavage. Treatment with either inhibitor alone or in combination showed a partial, non-significant effect on M2 cleavage (Figure S1E-F). Taken together, these results point to a dominant role for caspase-6 in M2 cleavage during IAV infection and a partial role for caspase-3. We conclude M2 is cleaved predominantly by caspase-6 with some redundancy, and decided to further elucidate the caspase cleavage site and its consequences during IAV infection.

A caspase cleavage motif has been predicted in the C-terminal motif of M2 [21]. Since caspases have a strict preference for aspartic acid at the scission bond, we mutated three amino acids from aspartic acid to alanine (D85A, D87A and D88A) to precisely identify the cleavage site [22]. We found that substitution of aspartic acid 87 or 88 with alanine (M2^D87A^ and M2^D88A^ mutants) had no effect on the appearance of the cleaved M2 product on immunoblots (Figure 1E-F). However, infection with IAV expressing M2^D85A^ abolished the lower molecular weight M2 band, indicating inhibition of caspase cleavage (Figure 1E-F, Figure S1G-H). To verify the tetrapeptide caspase-recognition motif is _82_SAVD_85_, we tested M2 cleavage in mutants including single amino acid changes upstream of D85 (M2^S82A^ and M2^V84A^). Both mutants were cleaved to a lesser extent than wild type M2, indicating these amino acids promote substrate recognition and cleavage (Figure 1E-F, Figure S1H). Together, these results show that during IAV infection, M2 is cleaved predominantly by caspase-6 and caspase-3, at the _82_SAVD↓A_86_ motif.

### M2 cleavage disrupts M2-LC3 interaction

We next asked whether cleavage of M2 at D85 had a functional consequence on the course of IAV infection. Downstream of the _82_SAVD↓A_86_ motif, an LC3 interacting region (LIR, aa 91-94) has previously been described [11] (Figure 1E). Therefore, we investigated whether M2 cleavage disrupted the M2-LC3 interaction.

Previous studies have reported that disruption of the M2 LIR motif reduces LC3B lipidation during infection [11]. To analyse the role of M2 cleavage, we assessed LC3B lipidation induced by M2 whilst caspases were inhibited by Z-VAD-FMK. LC3B lipidation was significantly increased upon with Z-VAD-FMK treatment, which correlated with the abrogation in the cleavage of M2. These observations were consistent with caspase-mediated disruption of the M2-LC3 interaction (Figure 2A-B).

**Figure 2.**
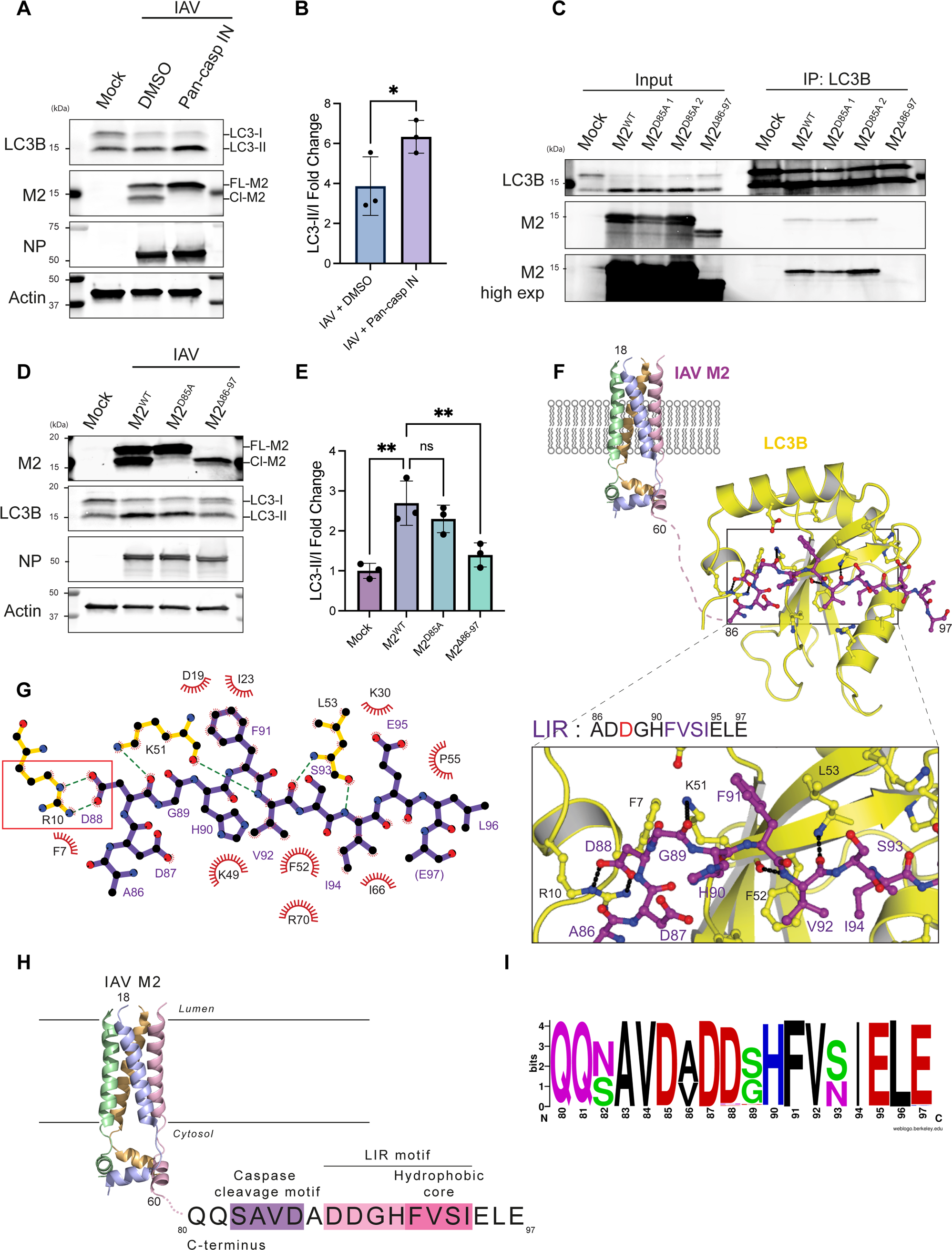
M2 cleavage disrupts M2-LC3 interaction. **A,** Representative immunoblots of lysates THP-1 cells infected with IAV PR8 for the indicated duration with an MOI of 10. After 8 hours, indicated samples were treated with DMSO as a control, or 20 µM Z-VAD-FMK as indicated for a further 16 hours. **B,** Quantification of **A**. Bars show mean ± SD of n = 3. *: P ≤ 0.05. Ordinary one-way ANOVA with Dunnett’s multiple comparisons. **C,** Immunoprecipitation of endogenous LC3B from A549 cells analysed by western blotting. Indicated samples were infected for 24 hours with IAV PR8 M2^WT^, or mutant strains, with an MOI of 10. **D,** Representative immunoblots of lysates of THP-1 cells infected with IAV PR8 M2^WT^, M2^D85A^, and M2^Δ86-97^ mutants for 24 hours with an MOI of 10. **E,** Quantification of **D**. Bars show mean ± SD of n = 3. **: P ≤ 0.01. Ordinary one-way ANOVA with Dunnett’s multiple comparisons. **F,** Model of IAV M2 proton channel (adapted from PDB 2RLF, aa 18-60) and LC3 in complex with M2 LIR. Crystal structure of LC3B in complex with M2 LIR: Ribbon and ball-and-stick representation model of the LC3B (Yellow) and M2 LIR (Purple, aa 86-97). **G,** LIGPLOT representation of the interaction between LC3B and M2 LIR. M2 LIR is shown in purple. The hydrogen bonds are in green dashed lines [48]. **H,** Diagram illustrating the IAV M2 proton channel (adapted from PDB 2RLF, aa 18-60). In the C-terminal region, the caspase cleavage motif (in purple) and preceded by the hydrophobic core (in pink) and acidic upstream amino acid (in light pink) of the LIR are highlighted. **I,** Conservation analysis of amino acids 80-97 in C-terminal region of M2. From 54974 human IAV M2 sequences, 2189 unique sequences were analysed [49]. Sequence logo was generated with WebLogo [50].

To verify the results obtained with pharmacological inhibitors, we next used a genetic approach to generate IAV expressing truncated M2 (M2^Δ86-97^) (Figure 1E, Figure S1G). We first assessed whether the M2^Δ86-97^ mutant, which lacks the LIR, was deficient in LC3B interaction through CoIP experiments (Figure 2C). These experiments showed that LC3B pulled down M2 during IAV M2^WT^ and M2^D85A^ infection. However, this interaction was not present during M2^Δ86-97^ infection (Figure 2C). This finding established that the M2^Δ86-97^ mutant is defective in LC3B interaction (Figure 2C). Concordant results were obtained in CoIP experiments performed in THP-1 cells (Figure S2A). Taken together, these results indicate that the M2-LC3 interaction is mediated through the LIR, and it is specifically disrupted by caspase cleavage.

We then assessed LC3B lipidation levels in THP-1 cells infected for 24 hours with IAV M2^WT^, M2^D85A^, and M2^Δ86-97^ mutants (Figure 2D-E). Consistent with M2 cleavage at D85, infection with the M2^D85A^ mutant only presented full-length M2 on immunoblots, while only the shorter M2 fragment was present in M2^Δ86-97^ infection (Figure 2D). We observed M2^WT^ and M2^D85A^ viruses induced comparable amounts of LC3B lipidation, while there was a significant decrease in LC3B lipidation induced by the M2^Δ86-97^ mutant (Figure 2D-E). This points to a crucial role of caspase-cleavage of M2 in modulating LC3B lipidation during infection. While LC3B lipidation during M2^Δ86-97^ mutant infection is significantly decreased, we reason that the residual LC3B lipidation is due to the proton channel activity of M2, which depends on the transmembrane region, and should be intact in distal C-terminal domain mutants (Figure 2D-E) [23].

To understand the interaction between M2, LC3B, and caspase cleavage at the molecular level, we obtained the co-crystal structure of the purified M2 C terminus (aa 70-97), containing the caspase cleavage site and LIR, and LC3B (aa 3-125). The structure was determined by molecular replacement using an LC3B structure (PDB: 2ZJD) as a search model (Figure 2F). One complex pair of M2-LC3 is in an asymmetric unit where F91 and I94 at position 1 and 4 of M2 LIR fit into the typical hydrophobic pockets of LC3B (Figure S2B). The region of M2 upstream from the LIR (amino acids 70-85), which contains the caspase cleavage motif, exhibited no clear electron density in our crystal structure due to its flexibility. The electron density of the four amino acids preceding the M2 LIR hydrophobic core was clearly observed (Figure 2G). We observed that the acidic side chain of D88 at −2 position of the M2 LIR forms two hydrogen bonds with the guanidinium group of R10 of LC3B, resulting in further stabilization of the M2-LC3 interaction (Figure 2G). Consistent with the observed structural role of D87 and D88 for binding, we observed a decrease in LC3B lipidation when THP-1 cells were infected with M2^D87A^ and M2^D88A^ (Figure S2C). Collectively, these data clarify the role of acidic amino acids in the C-terminal region of M2. While D85 lies in a flexible region of M2 consistent with its role in caspase cleavage, D87 and D88 are important for stabilising M2-LC3 binding.

These results point to consecutive linear motifs, with the _82_SAVD↓A_86_ motif cleaving the hydrophobic core of the LIR motif (aa 91-94) and acidic amino acids upstream of it, which we have identified as relevant for the M2-LC3B interaction (Figure 2H). Furthermore, analysis of 2189 unique M2 sequences in human IAV genomes showed high conservation of D85, consistent with this motif being important for viral fitness (Figure 2I).

### Cleaved M2 is impaired in plasma membrane relocalization

As M2 must travel to the plasma membrane for IAV budding, we assessed the impact of M2 cleavage on plasma membrane localisation of M2 [24]. To this end, we infected THP-1 cells with IAV M2^WT^, M2^D85A^, and M2^Δ86-97^ mutants, and performed M2 immunofluorescence with and without permeabilization to assess internal and surface M2 abundance, respectively. The M2^WT^ and M2^D85A^ mutants showed similar internal localisation, with a pool of M2 in the perinuclear region and substantial amounts at the plasma membrane (Figure 3A, Figure 3B). In M2^Δ86-97^ mutant infection, M2 was still present in the perinuclear region, but localised less to the plasma membrane (Figure 3A, Figure 3B).

**Figure 3.**
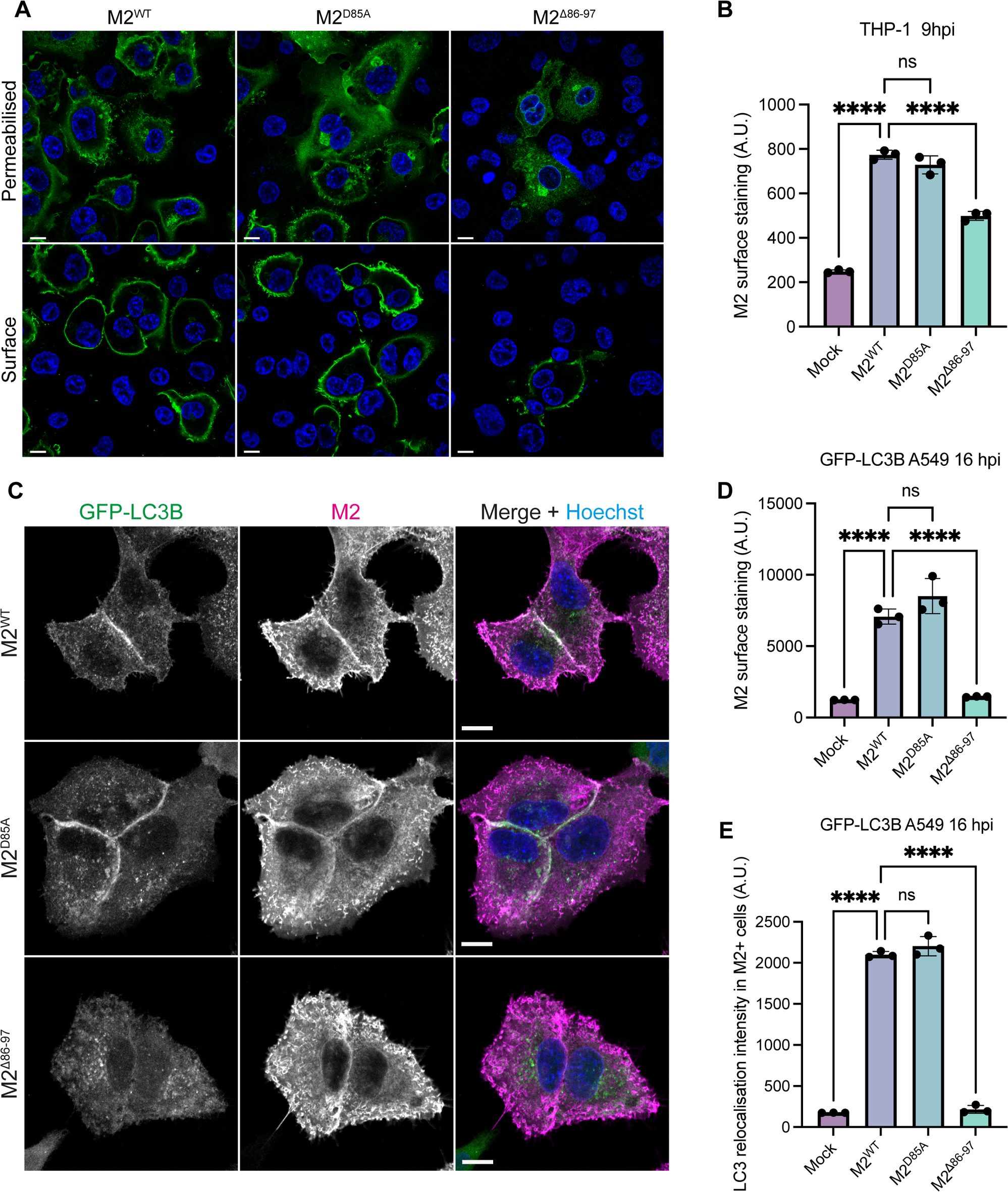
Cleaved M2 is impaired in plasma membrane relocalisation. **A,** Representative images of THP-1 cells infected IAV PR8 M2^WT^, M2^D85A^, and M2^Δ86-97^ mutants for 8 hours with an MOI of 1. Samples were stained with M2 (in green) and Hoechst (blue). Scale bar represents 10 µm. **B,** Quantification of M2 intensity in surface stained THP-1 cells. Samples were infected with IAV PR8 M2^WT^, M2^D85A^, and M2^Δ86-97^ for 8 hours with an MOI of 1. Bars show mean ± SD of n = 3. ****: P ≤ 0.0001. Ordinary one-way ANOVA with Dunnett’s multiple comparisons. **C,** Representative images of GFP-LC3B A549 cells infected with IAV PR8 M2^WT^, M2^D85A^, and M2^Δ86-97^ mutants for 16 hours with an MOI of 10. Images show GFP-LC3B (green), M2 (magenta), and Hoechst (blue). Scale bar represents 10 µm. **D,** Quantification of M2 intensity in surface stained GFP-LC3B A549 cells. Samples were infected with IAV PR8 M2^WT^, M2^D85A^, and M2^Δ86-97^ for 16 hours with an MOI of 10. Bars show mean ± SD of n = 3. ****: P ≤ 0.0001. Ordinary one-way ANOVA with Dunnett’s multiple comparisons. **E,** Quantification of GFP-LC3B spot intensity in M2 positive GFP-LC3B A549 cells. Samples were infected with IAV PR8 M2^WT^, M2^D85A^, and M2^Δ86-97^ for 16 hours with an MOI of 10. Bars show mean ± SD of n = 3. ****: P ≤ 0.0001. Ordinary one-way ANOVA with Dunnett’s multiple comparisons.

It has been reported that the M2 LIR affects LC3B localisation to the plasma membrane [11]. To answer whether cleavage of the LIR motif affected M2 trafficking, we infected GFP-LC3B A549 cells with IAV M2^WT^, M2^D85A^, and M2^Δ86-97^ mutants (Figure 3C). In permeabilised cells, M2 was localised in the cytoplasm and around the plasma membrane region during infection (Figure 3C). GFP-LC3B signal could be found colocalising with M2 during IAV M2^WT^ and M2^D85A^ infection, but not in M2^Δ86-97^ infection (Figure 3C). To recapitulate our results from THP-1 cells, GFP-LC3B A549 cells were surface stained with M2, and M2 intensity was quantified. We found M2^WT^ and M2^D85A^ mutants expressed similar amounts of M2 on the plasma membrane (Figure 3D). However, the M2^Δ86-97^ infection led to reduced amounts of M2 in the plasma membrane surface (Figure 3D). Furthermore, we assessed GFP-LC3B intensity in M2-positive cells, which confirmed that while M2^WT^ and M2^D85A^ infection induced a comparable amount of GFP-LC3B lipidation, GFP-LC3B spot intensity was significantly reduced in M2^Δ86-97^ infection (Figure 3E). These results suggest that the interaction of M2 with LC3B affects M2 transport to the plasma membrane.

### IAV M2^Δ86-97^ exhibits a decreased titer and reduced M2 incorporation into virions

To assess whether the decreased transport of M2 to the plasma membrane altered the composition and presence of different forms of M2 in virions, we purified IAV infectious particles from supernatant of IAV M2^WT^ infected MDCK cells at 48 hpi. Viral proteins could be identified at their expected molecular weights following Coomassie staining of purified infectious particles following SDS-PAGE (Figure 4A). Cell lysate and infectious particles were run in parallel to assess ratios of full-length to cleaved M2 present. While most M2 present in cells was the cleaved form, there was a 1:1 mix of full-length to truncated M2 in infectious particles (Figure 4B-C). This indicates that full-length M2 is preferentially incorporated into virions, which is consistent with its greater plasma membrane trafficking efficiency.

**Figure 4.**
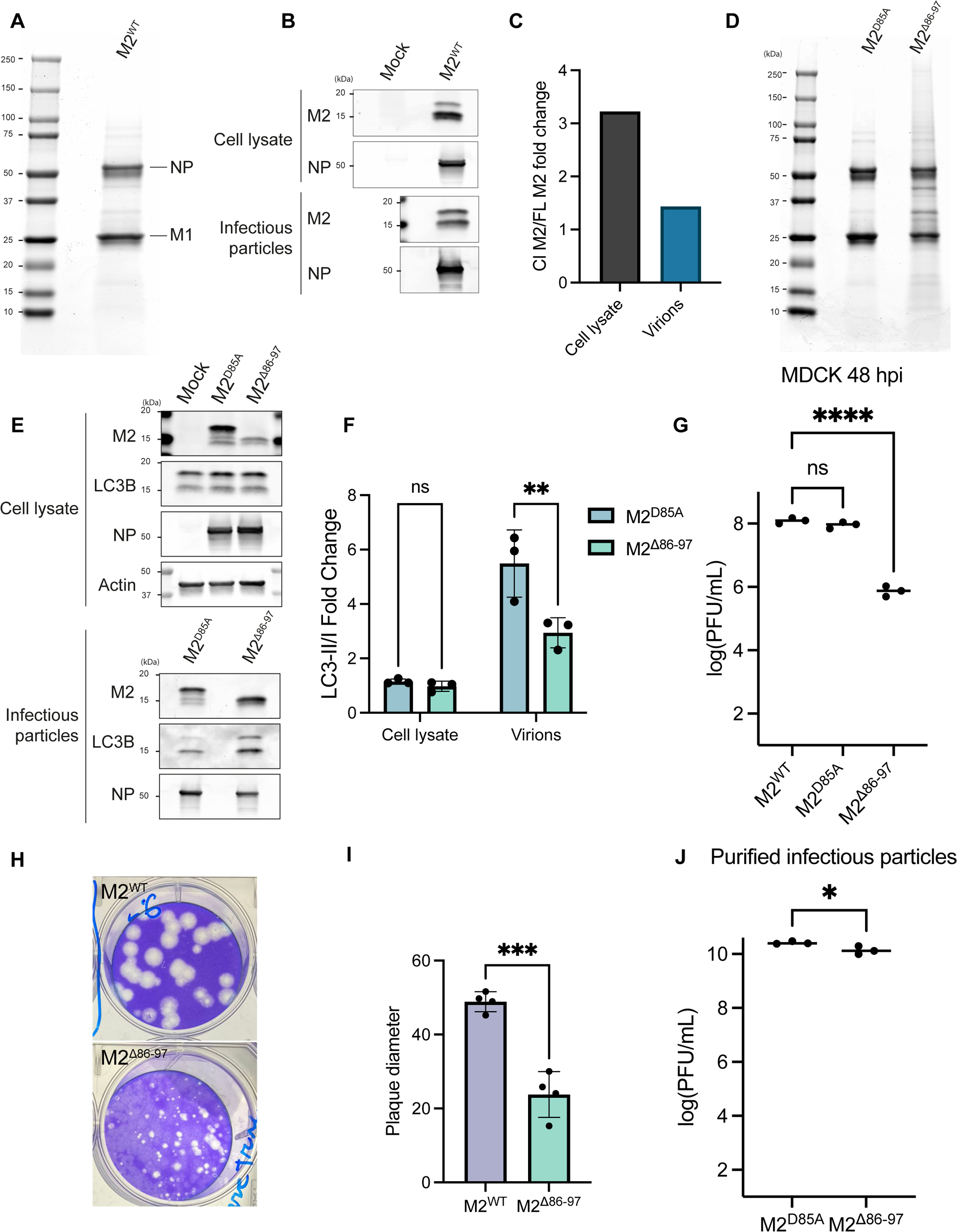
Cleaved M2 is incorporated into virions at a lower rate and exhibits a decreased titer. **A,** Coomassie stained gel of purified infectious particles from supernatant of MDCK cells infected for 48 hours with IAV PR8 M2^WT^. **B,** Representative immunoblots of cell lysate from MDCKs and infectious particles following purification. Cells were infected with IAV PR8 M2^WT^ for 48 hours. Supernatant was then collected and purified. **C,** Quantification of ratio of cleaved and full length M2 in MDCK cell lysate and purified virions, shown in **B**. **D,** Coomassie stained gel of purified infectious particles from supernatant of MDCK cells infected for 48 hours with IAV PR8 M2^D85A^, and M2^Δ86-97^ mutant as indicated. **E,** Representative immunoblots of cell lysate from MDCKs and infectious particles following purification. Cells were infected with IAV PR8 for 48 hours. Supernatant was then collected and purified. **F,** Quantification of **E**. Bars show mean ± SD of n = 3. **: P ≤ 0.01. Two-way ANOVA with Šidák multiple comparisons test. **G,** Plaque assay quantification to assess IAV titer. Supernatant of MDCKs infected with IAV PR8 M2^WT^, M2^D85A^, and M2^Δ86-97^ mutants was collected after 48 hours. Plaque assays were performed for 48 hours. Bars show mean ± SD of n = 3. ****: P ≤ 0.0001. Log-transformed data are plotted. Ordinary one-way ANOVA with Dunnett’s multiple comparisons. **H,** Images of plaque assay plates stained with toluidine blue. **I,** Quantification of plaque diameter from **H**. Bars show mean ± SD of n = 4. ***: P ≤ 0.001. Unpaired t test. **J,** Plaque assay quantification to assess IAV infectious particle plaque forming units/mg of purified protein to assess titer. Purified infectious particles of M2^D85A^ or M2^Δ86-97^ mutants were used. Plaque assays were performed for 48 hours. Bars show mean ± SD of n = 3. *: P ≤ 0.05. Graph shows data as Y=log(Y). Unpaired t test.

We then proceeded to purify infectious particles from supernatant of M2^D85A^ or M2^Δ86-97^ infected MDCKs at 48 hpi (Figure 4D). Coomassie staining of purified virions following SDS-PAGE suggested both viral mutants were broadly comparable, with the main IAV proteins identified (Figure 4D). We observed that the ratio of LC3II/LC3I in virions was reduced in M2^Δ86-97^ viral particles when compared to M2^D85A^, which indicates more lipidated LC3B was incorporated into virions containing the M2 LIR motif (Figure 4E-F). We next performed proteomic analysis of purified M2^D85A^ and M2^Δ86-97^ virions, which showed that LC3B was more abundant in M2^D85A^ virions (Figure S3A). Furthermore, the ratio of viral proteins to M1 abundance was similar between the M2^D85A^ and M2^Δ86-97^ mutants. However, the overall abundance of viral proteins was significantly increased in M2^D85A^ purified infectious particles (Figure S3A-B). This is consistent with a deficiency in budding present in the M2^Δ86-97^ mutant.

To assess whether the introduction of uncleavable or truncated M2 in IAV resulted in attenuated virus, we performed plaque assays to assess the titer (Plaque Forming Units (PFU)/mL) formed by each mutant. While M2^WT^ and M2^D85A^ had comparable titers, the M2^Δ86-97^ mutant had a significantly decreased titer (Figure 4G). This indicates that caspase cleavage of M2 produces an attenuated virus. Consistent with this, the plaque size produced by M2^Δ86-97^ infection was significantly decreased when compared to plaques formed by M2^WT^ infection (Figure 4H-I). To understand the stage of viral life cycle affected by cleavage, we infected THP-1 cells for 8 hours with IAV M2^WT^, M2^D85A^, and M2^Δ86-97^ mutants. We performed a plaque assay of the supernatant to assess titer, which revealed that the M2^Δ86-97^ mutant titer was still significantly decreased (Figure S3C). Purified M2^D85A^ and M2^Δ86-97^ infectious particles were used to perform a plaque assay and quantify titer (Figure 4J). Here, we could observe the difference in titer between M2^D85A^ and M2^Δ86-97^ mutants was reduced when compared to previous titers (Figure 4G, 4J, S3C). Taken together, these results suggest a defect in assembly or budding of IAV with M2^Δ86-97^.

## Discussion

Here, we have defined a caspase cleavage site in IAV M2 by mutagenesis (_82_SAVD_85_). Furthermore, we have solved a structure of the immediately adjacent LIR motif in the otherwise unstructured IAV M2 C-terminal in complex with LC3B. Mutagenesis of D87 and D88 acidic residues confirmed their contribution to the LIR motif, while having no effect on caspase cleavage. Therefore, the concatenation of caspase cleavage site and LIR suggests that two consecutive protein motifs act together to alter M2 function. This parallels a previous description of caspase-mediated cleavage of the autophagy receptor p62, directly upstream of a LIR motif [25]. We conclude that this highly conserved interaction and cleavage act as a regulatory mechanism exploited by IAV to fine-tune virion production in different cellular contexts.

Caspases require cleavage motifs to be accessible, being frequently observed in unstructured regions of proteins [26, 27]. This is consistent with the flexible nature of the caspase cleavage motif observed in the structure of the M2-LC3B interactions. Using CRISPR generated caspase deficient cell lines and chemical inhibition, we show that caspase-6 is largely responsible for M2 cleavage. A partial reduction in M2 cleavage was observed with caspase-3 deficiency and inhibition. This effect may be indirect, since caspase-3 has been reported to cleave caspase-6 for activation [28]. The possibility also remains that, while executioner caspases perform distinct roles during apoptosis, they may exhibit a degree of redundancy when it comes to viral substrates [29].

Plasma membrane localisation of LC3 has previously been reported during IAV infection. This relocalisation is abolished during infection with LIR deficient virus [11]. Here, we show that M2 and LC3 transport to the plasma membrane can be regulated by caspase-mediated removal of the LIR motif, contextualising previous findings. IAV uses the host plasma membrane to form its own envelope [30]. The C-terminal region of M2 is crucial for viral assembly and budding [31, 32]. Here, we show for the first time inclusion of LC3 into IAV virions through interaction with M2. The inclusion of LC3 in virions has been reported in other viruses, although the functional importance for this is unclear [33–37].

The high conservation level of caspase cleavage and LIR motifs highlights the complexity of host machinery regulation by IAV. Caspase cleavage of M2 must provide an evolutionary advantage for the virus despite attenuation observed in the permanently truncated M2^Δ86-97^ mutant. It has been previously demonstrated that the LIR motif is required for optimal filamentous budding [11]. Pathogenic isolates of IAV are pleiomorphic, exhibiting both filamentous and spherical virions [38]. While the exact role of viral filament formation is unknown, it may aid infection of neighbouring cells in the upper respiratory tract [38, 39]. Filament budding appears to be inefficient, using more host plasma membrane for budding than spherical virions [38]. Therefore, we speculate that caspase cleavage of M2 removes the LIR motif to enable a switch between filamentous and non-filamentous budding in response to depletion of cellular resources. This would provide an evolutionary advantage to the virus, maximising virion production at later time points.

## Limitations

We hypothesise that caspase cleavage of M2 may have a role in viral budding, and may impact the ability of IAV to form filaments. However, we were unable to rescue an IAV M2^Δ86-97^ virus in a filamentous background.

Our work reports the importance of caspase cleavage in modulating LC3B lipidation, trafficking to the plasma membrane, and inclusion into virions. We do not exclude that this may be occurring with other Atg8 proteins.

## Acknowledgements

The authors thank the following: Lewis Timimi, Rachel Ulferts, and other past and present members of the Beale and Shenoy labs for helpful comments and suggestions. We thank E. Hutchinson for helpful advice on influenza virion purification. We thank the Proteomics, High Throughput Screening, Cell Services and Light Microscopy core facilities (Francis Crick Institute) for support.

CFN, BM, MJ, CD and RB were supported by The Francis Crick Institute, which receives its core funding from Cancer Research UK and the Medical Research Council. This research was funded in whole, or in part, by the Wellcome Trust. AS acknowledges funding from the MRC (MR/T00004X/1). For the purpose of Open Access, the author has applied a CC BY public copyright licence to any Author Accepted Manuscript version arising from this submission.

## Author Contributions

CFN, AS and RB designed the study. CFN performed all experiments and analysis except as listed below. MA and DM solved the LC3B/M2 crystal structure. BM generated cell lines. MJ performed high throughput microscopy analysis. CD performed proteomics experiments. AS and RB provided supervision and oversight. CFN and RB wrote the original manuscript, and all authors contributed to improving the manuscript.

## Methods

### Antibodies, reagents, and chemicals

Antibodies used in this study were mouse monoclonal anti-IAV Matrix 2 (Abcam, ab5416, WB 1:1000, IF 1:100), rabbit monoclonal anti-LC3B (Novus Biologicals, NBP2-46892, WB 1:1000), rabbit polyclonal caspase-6 (Cell Signaling, 9762, WB 1:1000), mouse monoclonal anti-β-actin (Proteintech, 66009-1, WB 1:10000), rabbit polyclonal anti-IAV NP (GeneTex, GTX125989, WB 1:10000), rabbit polyclonal PARP-1 (Cell Signaling, 9542, WB 1:1000).

Chemical reagents used in this study include Z-VAD-FMK (InvivoGen; tlrl-vad), Z-VEID-FMK (R&D systems; FMK006), Z-DEVD-FMK (R&D systems FMK004), and Z-IETD (Invivogen; inh-ietd) at indicated concentrations. Ammonium bicarbonate, phosphoric acid, triethylammonium bicarbonate (TEAB), and sodium dodecyl sulfate (SDS) were purchased from Sigma-Aldrich (St. Louis, MO). Formic acid (FA) 99 %, iodoacetamide (IAA), tris(2-carboxyethyl)phosphine hydrochloride (TCEP), trypsin protease MS-grade were purchased from Thermo Scientific Inc. (Rockford, IL, USA). LCMS-grade solvents: water, acetonitrile (ACN), methanol, were purchased from Fisher Chemical. S-Trap micro columns were purchased from ProtiFi (Huntington, NY). Evotips were purchased from Evosep (Odense, Danemark).

### Cell culture

THP-1 cells (provided by the Cell Services STP of the Francis Crick Institute) were grown in Roswell Park Memorial Institute (RPMI; GIBCO Life Technologies) containing 10 % Fetal Calf Serum (FCS), GlutaMax (GIBCO Life Technologies) and Penicillin-Streptomycin (GIBCO Life Technologies). For differentiation, THP-1 cells were treated with 100 ng/mL PMA for 48 hours. Media was changed and cells were incubated in fresh media for a further 24 hours. HEK293T, MDCK, and A549 cells (provided by the Cell Services STP of the Francis Crick Institute) were cultured in Dulbecco’s modified Eagle’s medium (DMEM; GIBCO Life Technologies) containing 10 % FCS, GlutaMax and Penicillin-Streptomycin. HAP1 cells, including Parental, CASP3 KO (ID: HZGHC001508c004), CASP6 KO (ID: HZGHC001509c004), CASP7 KO (ID: HZGHC001510c003), and CASP8 KO (ID: HZGHC001511c007) cells were purchased from Horizon Discovery, and grown in Iscove’s modified Dulbecco’s medium (IMDM; GIBCO Life Technologies) containing 10 % FCS and Penicillin-Streptomycin. All cell lines were grown in incubators at 37 °C in 5 % CO_2_.

### Plasmids

Plasmids for IAV generation through reverse genetics are described in [40]. pENTR-Flag-CASP6 was produced using the pENTR/D-TOPO Cloning Kit (Invitrogen) with the CASP6_OHu19575_pcDNA3.1(+)-N-DYK plasmid. pLenti-PGK-Flag-CASP6-HygR was generated through gateway cloning with LR clonase (Invitrogen), with pENTR-Flag-CASP6 and pLenti-PGK-GWT-HygR.

### Lentivirus generation

pLenti-CASP6_OHu19575_pcDNA3.1(+)-N-DYK plasmid was used to generate lentivirus. Lentivirus was produced by co-transfection of psPAX2 and pMD2-G with PEI in HEK293T cells. Cells were transduced through spinoculation with lentivirus and 8 µg/mL polybrene for 1 hour at 500 x *g* at room temperature. Transduced cells were selected with 200 µg/mL Hygromycin.

### SDS-PAGE and Western blot

Cells were grown, treated, or infected in 6, 12 or 48 well plates. Before lysis, cells were washed twice in PBS. Cells were then lysed in ice-cold NP-40 buffer (0.5 % NP-40, 25 mM Tris–HCl (pH 7.5), 100 mM NaCl, 50 mM NaF) with fresh protease inhibitor cocktail (Sigma). Cell lysate was then collected, clarified (16,200 x *g*, 4 °C, for 30 minutes) and the sample protein concentration was assessed through a BCA assay (Pierce). Proteins were resolved on Mini-PROTEAN®TGX gels (Bio-Rad). Staining was carried out with InstantBlue® Coomassie Protein Stain (Abcam) or transferred onto nitrocellulose membranes for Western blot.

Nitrocellulose membranes were blocked in 5 % dry milk powder in 0.1 % Tween-20 for 1 hour at room temperature. Primary antibodies were incubated at room temperature for 1 hour, or overnight at 4 °C. Membranes were washed in 0.1 % Tween-20 and then stained with species-specific IRdye 800CW and 680LT coupled secondary antibodies (LI-COR). Membranes were washed and scanned with an Odyssey CLx scanner (LI-COR).

### Immunofluorescence

Cells were grown in coverslips or in clear, flat-bottom 96-well (PerkinElmer, 6055300), and then fixed with a final concentration of 4 % formaldehyde in PBS for 20 minutes. For samples requiring permeabilization, cells were treated with 0.2 % Triton-X100 for 5 minutes, and washed with PBS. All samples were blocked in 3 % BSA in PBS for 30 minutes. Samples were incubated with primary antibodies for 1 hour, washed, and incubated with species-specific AlexaFluor 488 and 647 (Thermo), and Hoescht 33342 for 45 minutes. Coverslips were mounted onto slides using ProLong™ Glass Antifade Mountant (Invitrogen).

Imaging was done with a Zeiss Invert880 (63X oil-immersion objective) or a Visitech iSIM 827 microscope (100x oil-immersion objective). Images were processed in Fiji 828 (v.2.3.0/1.53f; Schindelin et al., 2012) and Adobe Photoshop 2023 (v.24.7.0).

### High-content immunofluorescence quantification

For quantifying the M2 protein, the cells were cultured and infected on 96well CellCarrier plates (PerkinElmer, 6005430). Plates were then scanned using Opera Phenix plus (PerkinElmer) after staining. The images were acquired utilising a 40x_water_NA1.1 lens with confocal of 8 planes Z-stacks spanning from −1 to 2.5 µM. The process involved the utilization of excitation lasers at wavelengths of 375, 488 nm, coupled with emission filters at 435-480 and 500-550. The quantification of M2 (488) was analysed using Harmony 5.0.

### Influenza A virus production and infection

Influenza A virus PR8 (strain A/Puerto Rico/8/1934) was generated through an eight plasmid-based system in HEK293T cells, as previously described [40]. Stocks were passaged in MDCK cells, in presence of TPCK-trypsin (Worthington) and 0.14 % BSA Fraction V. For infection, cells were first washed with serum-free media, and then incubated with virus in serum-free media at 37 °C for an hour. Inoculum was then replaced with fully complemented media and incubated at 37 °C for the indicated period.

### Reverse genetics

Influenza A virus PR8 (strain A/Puerto Rico/8/1934) mutants were generated through the above eight plasmid-based system [40]. Mutant segment 7 plasmids were generated through Q5 Site-Directed Mutagenesis Kit (New England BioLabs).

### Immunoprecipitation

Cells were incubated in lysis buffer (0.5 % NP-40, 10 mM Tris–HCl (pH 7.5), 150 mM NaCl, 0.5 mM EDTA) at 4 °C rotating for 30 minutes, and clarified (16,200 x *g*, 4 °C, for 30 minutes). Dilution buffer (10 mM Tris–HCl (pH 7.5), 150 mM NaCl, 0.5 mM EDTA) was added to lysed samples. Dynabeads™ Protein G for Immunoprecipitation (ThermoFisher 10003D) were washed with PBS-Tween, and incubated with relevant antibodies for 30 minutes at 4 °C. Samples were then incubated with beads for 2 hours rotating at 4 °C. Samples were washed 5 times, resuspended in sample buffer, and boiled for 5 minutes at 95 °C before running.

### Crystallization, X-ray data collection and structure determination

M2 LIR (70-97) and LC3B (3-125) fusion protein in pETM60 were expressed as N-terminal Ubiquitin His tag fusion proteins in *Escherichia coli* BL21 (DE3) [41]. Cells were grown at 37 °C to an OD_600_ of 0.5, followed by induction with 0.5 mM isopropyl-β-d-thiogalactoside and further incubation at 20 °C for 20 hours. Cells were lysed by sonication in lysis buffer (25 mM Tris, 200 mM NaCl, pH 8.5), and the lysate was cleared by centrifugation (15000 x *g*, 4 °C, 40 minutes). The expressed protein was purified by TALON Metal Affinity Resin (Clontech), cleaved by TEV protease, and purified by size exclusion chromatography (HiLoad 16/600 Superdex 75 column, GE Life Sciences) in 25 mM Tris, 200 mM NaCl, pH 8.5. The crystals of LC3B in complex with M2 LIR were obtained using 30 % Jeffamine, 0.1 M HEPES, pH 7.0 by the sitting-drop vapor diffusion method at 293K. Diffraction data were collected at Swiss Lightsource SLS, beam line PXIII and processed with XDS [42]. The crystal structure was determined by molecular replacement using the LC3B structure (PDB: 2ZJD) as a search model. Manual model building and refinement were done with Coot, CCP4 software suite and Phenix [43–45]. The final statistics of refined models are shown in Supplementary Table S1, and the atomic coordinates have been deposited in the Protein Data Bank (PDB ID: 8YV6).

### Plaque assays

To quantify viral titers, virus was serially diluted in serum free media. 200 µL of diluted virus at different dilutions were added to PBS-washed MDCK cells. Cells were incubated at 37 °C for an hour, with shaking every 15 minutes. Cells were then overlaid with serum free media, supplemented with 50 % Avicel (RC-581, IMCD), TPCK-trypsin (Worthington) and 0.14 % BSA Fraction V, and incubated for 48 hours. Supernatant was then removed, and cells were fixed with Neutral Buffered Formalin for 20 minutes. Staining was performed with toluidine blue for 20 minutes.

### Viral purification

Viral purification was performed according to [46]. MDCK supernatants infected with relevant viruses were spun at 4 °C for 10 minutes at roughly 2000 x *g*, in a table-top centrifuge. Media was then transferred to thickwall ultracentrifuge tubes (Beckman Coulter, 355642), placed SW28 rotor, and spun for 10 minutes at 4 °C at 18,000 x *g*. The supernatant was gentle layered on top of 5 mL of chilled 30 % sucrose. Virions were pelleted for 90 minutes at 4 °C at 112,000 x *g*. Supernatants were removed, and the pellet was resuspended in 50 µL of 4 °C PBS, and pooled. Resuspended pellets were layered on top of a sucrose density gradient (30-60 %) prepared in thinwall ultracentrifuge tubes (Beckman Coulter, 331372), which were placed in SW41 rotor buckets, and spun 150 minutes at 4 °C at 209,000 x *g*. A milky band was visible at 40 % sucrose. The band was collected with a Pasteur pipette, and ejected into a tube with 9 mL of 4 °C PBS. Finally, virions were pelleted for 60 minutes at 4 °C at 154,000 x *g*. The supernatant was discarded, and remaining pellet was resuspended in 80 µL of 4 °C PBS.

### Proteomic sample preparation

Protein digestion in the S-Trap micro column was performed according to the manufacturer’s protocol with some modifications. Briefly, 100 µg protein in lysis buffer (10 % SDS in 100 mM TEAB pH 8.5) was reduced by adding TCEP to the protein solution to a final concentration of 5 mM and incubated for 15 minutes at 55 °C. The alkylation was then performed by adding IAA to the reduced samples to a final concentration of 20 mM and incubated for 30 minutes at room temperature in the dark. A final concentration of 1.1 % (v/v) phosphoric acid followed by six-fold volume of binding buffer (90 %, v/v, methanol in 100 mM TEAB) was next added to the protein solution. After vortexing, the solution was loaded into an S-Trap micro column. The solution was removed by spinning the column at 2,000 x *g* for 1 min. The column was washed with 400 µL binding buffer three times. Finally, 125 µL of 0.80 mg/mL trypsin in 50 mM ammonium bicarbonate was added to the column and incubated 18 hours at 37 °C. Digested peptides were eluted using 80 µL of three buffers consecutively: (1) 50 mM ammonium bicarbonate, (2) 0.2 % (v/v) FA in H_2_O, and (3) 50 % (v/v) ACN. The elution solutions were collected in the same tube and dried under vacuum.

### Proteomic sample acquisition DIA

Dried samples were resuspended in 0.1 % (v/v) FA and 200 ng of tryptic peptides were loaded onto Evotip and analysed using an Evosep One LC (EVOSEP) connected to a timsTOF Pro (Bruker). The Evosep One method was 60 SPD (21-minute gradient, cycle time of 24 minutes) and the mass spectrometer was operated in DIA-PASEF mode. The DIA-PASEF method consisted of twelve *m/z* windows and a cycle time of 1.37 s. The other mass spectrometer parameters were set as follow: *m/z* range, 262 to 1200, the mobility (1/K0) range was set to 0.60 to 1.60 V.s/cm^2^, and the accumulation and ramp time were 100 ms.

### Proteomic sample search

The viral proteins from IAV strain A/Puerto Rico/8/1934 (Uniprot Taxon ID 211044) and host proteins from Canis lupus familiaris (Madin-Darby canine kidney cell line) (Uniprot Taxon ID 9615) FASTA files from the Uniprot database were used to generate a spectral library within DIA-NN 1.8 [47] (Data-Independent Acquisition by Neural Networks) following a library-free search method. Specific search settings included cysteine carbamidomethylation enabled as a fixed modification, N-terminal methionine excision enabled, maximum missed cleavages 1, min precursor + 2, max precursor + 4, neural network classifier was set to single-pass mode, cross run normalisation was set to retention time dependent and library generation was set to smart profiling. All output was filtered at 0.01 false discovery rate (FDR).

### Statistical analysis

Statistical analysis was performed in GraphPad Prism 10 Version 10.1.1 (270). Statistical tests used, error bars, number of replicates, and statistical significance are reported in figure legends.

---

**Figure S1.**
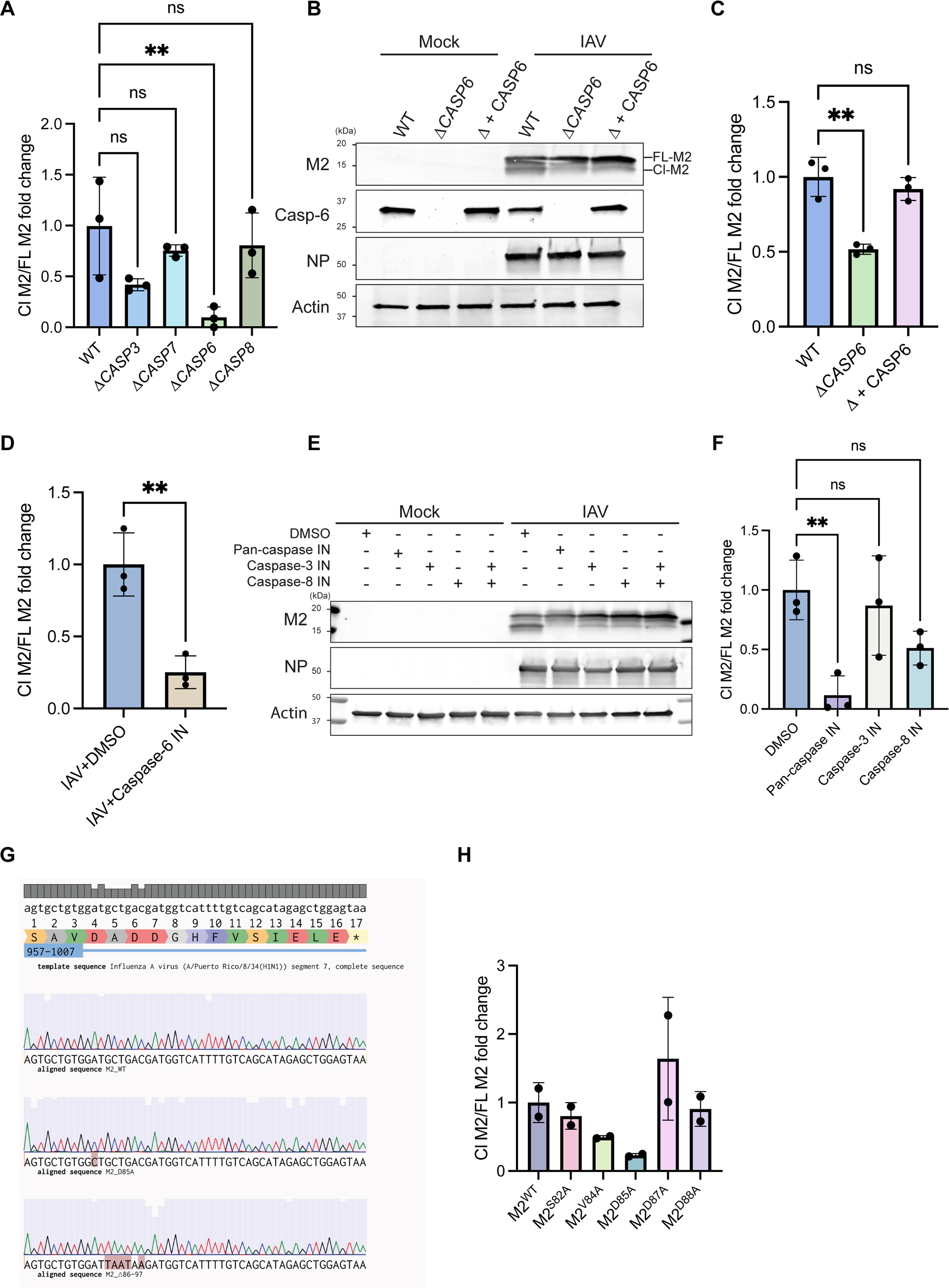
Extended data supporting “M2 is cleaved at SAVD motif by caspases”. **A,** Quantification of Figure 1C. Bars show mean ± SD of n = 3. **: P ≤ 0.01. Ordinary one-way ANOVA with Bartlett’s multiple comparisons. **B,** Representative immunoblots of lysates of wild type (WT), Δ*CASP6*, and Δ*CASP6* stably expressing Caspase-6. Indicated samples were infected for 16 hrs with IAV PR8 at an MOI of 10. **C,** Quantification of **B**. Bars show mean ± SD of n = 3. **: P ≤ 0.01. Ordinary one-way ANOVA with Dunnett’s multiple comparisons. **D,** Quantification of Figure 1D. Bars show mean ± SD of n = 3. **: P ≤ 0.01. Unpaired t test. **E,** Representative immunoblots of lysates of THP-1 cells infected with IAV PR8 for 24 hours. Indicated samples were treated with DMSO as a control, or 50µM pan-caspase inhibitor (Z-VAD-FMK), caspase-3 inhibitor (Z-DEVD-FMK), and caspase-8 inhibitor (Z-IETD-FMK) as indicated. **F,** Quantification of **E**. Bars show mean ± SD of n = 3. **: P ≤ 0.01. Ordinary one-way ANOVA with Dunnett’s multiple comparisons. **G,** Multiple sequence alignment of amino acids 82-97 in IAV M2. IAV PR8 M2^WT^, M2^D85A^, and M2^Δ86-97^ sequences were produced with Sanger sequencing and aligned to Influenza A virus (A/Puerto Rico/8/34(H1N1)) segment 7 [51]. **H,** Quantification of Figure 1F. Bars show mean ± SD of n = 2.

**Figure S2.**
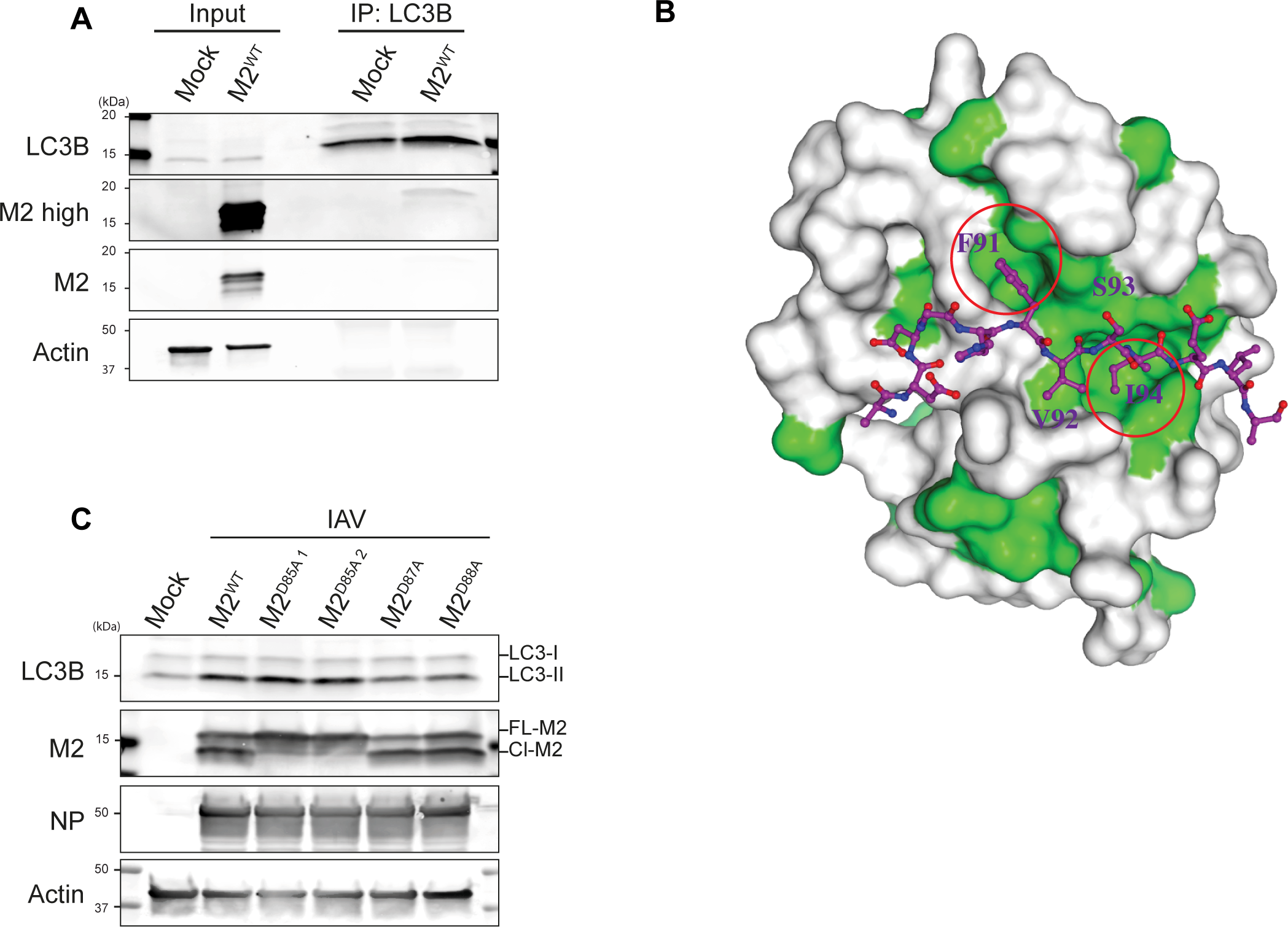
Extended data supporting “M2 cleavage disrupts M2-LC3 interaction”. **A,** Immunoprecipitation of endogenous LC3B from A549 cells analysed by western blotting. Indicated samples were infected for 24 hours with IAV PR8 WT, or mutant strains, with an MOI of 10. **B,** Surface representation model of M2-LC3B LIR complex. Hydrophobic residues of LC3B are coloured green. Red circles indicate hydrophobic pockets of LC3B for LIR interaction. **C,** Representative immunoblots of lysates of THP-1 cells infected with IAV M2^WT^, M2^D85A^, M2^D87A^and M2^D88A^ mutants for 24 hours with an MOI of 10.

**Figure S3.**
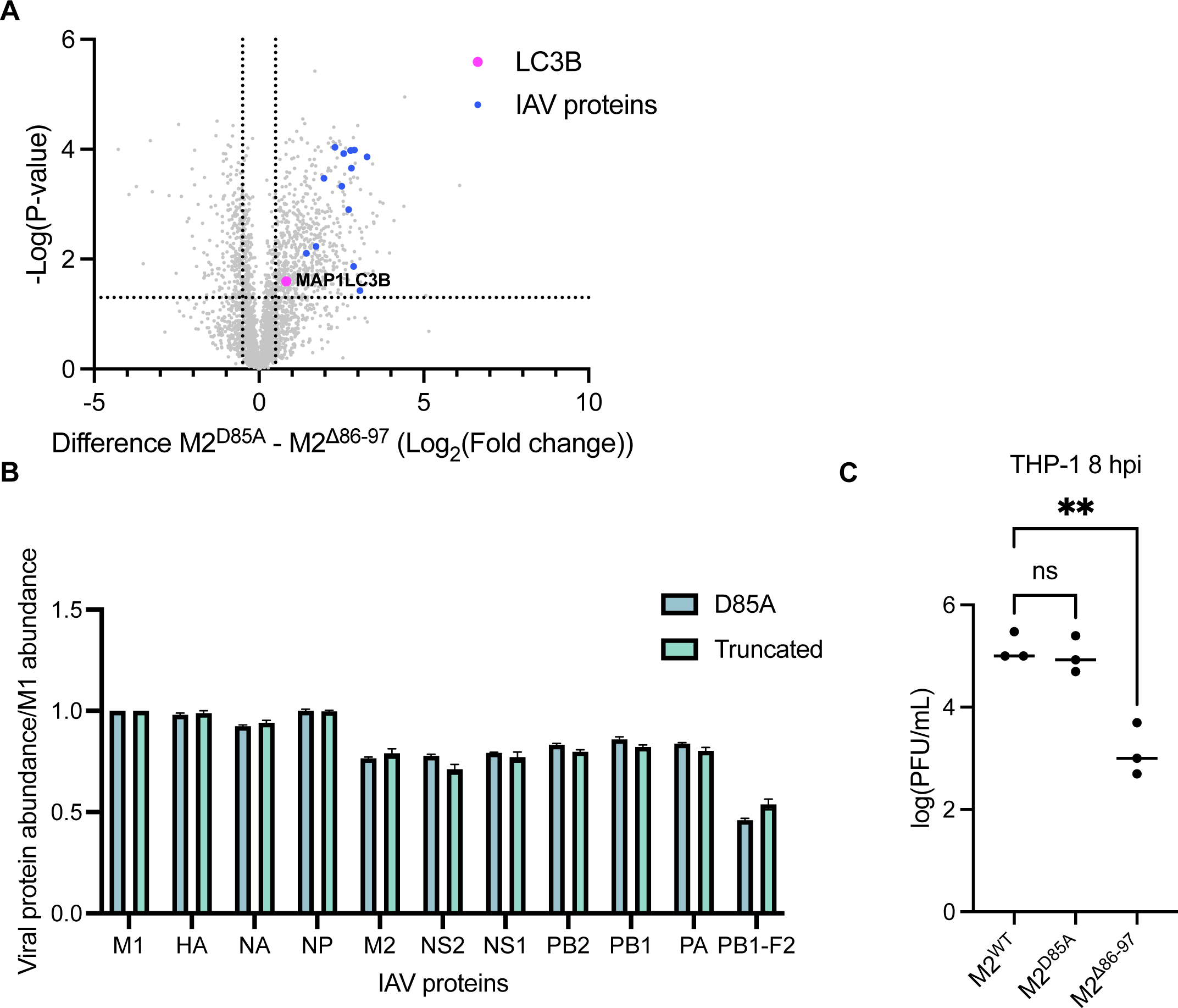
Extended data supporting “Cleaved M2 is incorporated into virions at a lower rate and exhibits a decreased titer”. **A,** Volcano plot showing the difference in log_2_(fold change) of abundance of proteins expressed in M2^D85A^ purified virions when comparing them to M2^Δ86-97^ purified virions. MAP1LC3B is shown as a pink dot, and Influenza A proteins are shown as blue dots. Dashed lines represent a difference in log_2_(fold change) of more than 0.5 or less than −0.5, and -Log(P-value) of 1.3). **B,** Abundance of viral proteins was calculated through normalisation to M1 abundance and compared between M2^D85A^ and M2^Δ86-97^ abundance. **C,** Plaque assay quantification to assess IAV titer following THP-1 infection. Supernatant of THP-1 cells infected with IAV PR8 M2^WT^, M2^D85A^, and M2^Δ86-97^ mutants was collected after 8 hours. Plaque assays were performed for 48 hours. Bars show mean ± SD of n = 3. **: P ≤ 0.01. Graph shows data as Y=log(Y). Ordinary one-way ANOVA with Dunnett’s multiple comparisons.

**Supplementary table 1.**
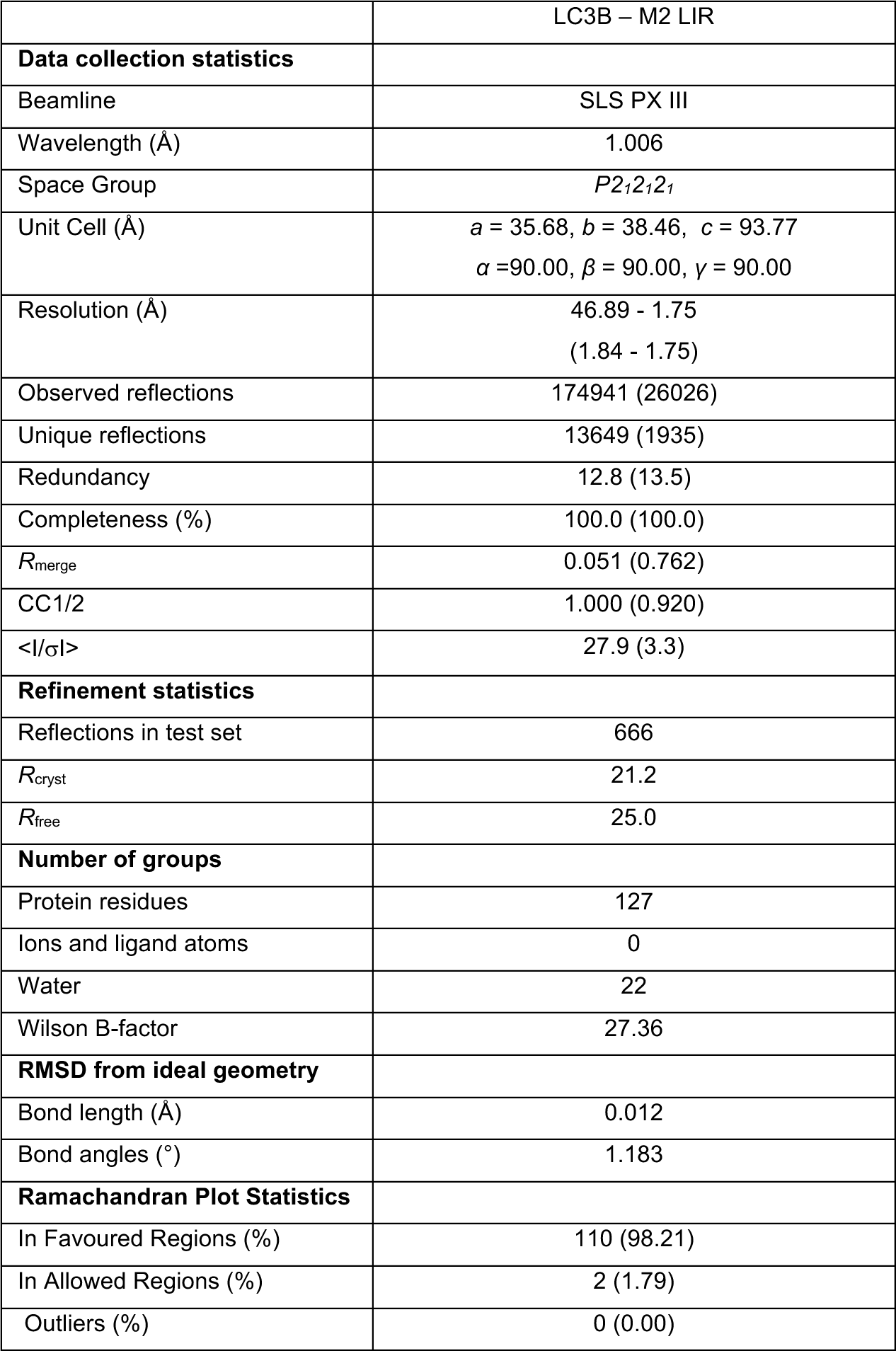
Crystallisation data collection and refinement statistics. Values in parentheses are for the highest resolution shell.

## References

1. WHO. Influenza (Seasonal) Fact Sheet. 2023 [cited 2024 04.03]; Available from: https://www.who.int/en/news-room/fact-sheets/detail/influenza-(seasonal).

2. Taubenberger, J.K. and J.C. Kash, Influenza virus evolution, host adaptation, and pandemic formation. Cell Host Microbe, 2010. 7(6): p. 440–51.

3. Lamb, R.A., Influenza, in Encyclopedia of Virology (Third Edition), B.W.J. Mahy and M.H.V. Van Regenmortel, Editors. 2008, Academic Press: Oxford. p. 95–104.

4. Holsinger, L.J., et al., Influenza A virus M2 ion channel protein: a structure-function analysis. J Virol, 1994. 68(3): p. 1551–63.

5. Pinto, L.H., L.J. Holsinger, and R.A. Lamb, Influenza virus M2 protein has ion channel activity. Cell, 1992. 69(3): p. 517–28.

6. Shimbo, K., et al., Ion selectivity and activation of the M2 ion channel of influenza virus. Biophys J, 1996. 70(3): p. 1335–46.

7. Scott, C., et al., Site-directed M2 proton channel inhibitors enable synergistic combination therapy for rimantadine-resistant pandemic influenza. PLoS Pathog, 2020. 16(8): p. e1008716.

8. Takeuchi, K. and R.A. Lamb, Influenza virus M2 protein ion channel activity stabilizes the native form of fowl plague virus hemagglutinin during intracellular transport. J Virol, 1994. 68(2): p. 911–9.

9. Alvarado-Facundo, E., et al., Influenza virus M2 protein ion channel activity helps to maintain pandemic 2009 H1N1 virus hemagglutinin fusion competence during transport to the cell surface. J Virol, 2015. 89(4): p. 1975–85.

10. Ciampor, F., et al., Evidence that the amantadine-induced, M2-mediated conversion of influenza A virus hemagglutinin to the low pH conformation occurs in an acidic trans Golgi compartment. Virology, 1992. 188(1): p. 14–24.

11. Beale, R., et al., A LC3-interacting motif in the influenza A virus M2 protein is required to subvert autophagy and maintain virion stability. Cell Host Microbe, 2014. 15(2): p. 239–47.

12. Ulferts, R., et al., Subtractive CRISPR screen identifies the ATG16L1/vacuolar ATPase axis as required for non-canonical LC3 lipidation. Cell Rep, 2021. 37(4): p. 109899.

13. Fletcher, K., et al., The WD40 domain of ATG16L1 is required for its non-canonical role in lipidation of LC3 at single membranes. The EMBO journal, 2018. 37(4): p. e97840.

14. Xu, Y., et al., A Bacterial Effector Reveals the V-ATPase-ATG16L1 Axis that Initiates Xenophagy. Cell, 2019. 178(3): p. 552–566.e20.

15. Timimi, L., et al., The V-ATPase/ATG16L1 axis is controlled by the V1H subunit. bioRxiv, 2023: p. 2023.12.19.572309.

16. Thornberry, N.A. and Y. Lazebnik, Caspases: enemies within. Science, 1998. 281(5381): p. 1312–6.

17. Puccini, J. and S. Kumar, Caspases, in Encyclopedia of Cell Biology, R.A. Bradshaw and P.D. Stahl, Editors. 2016, Academic Press: Waltham. p. 364–373.

18. Richard, A. and D. Tulasne, Caspase cleavage of viral proteins, another way for viruses to make the best of apoptosis. Cell death & disease, 2012. 3(3): p. e277–e277.

19. Zhirnov, O.P. and H.-D. Klenk, Alterations in caspase cleavage motifs of NP and M2 proteins attenuate virulence of a highly pathogenic avian influenza virus. Virology, 2009. 394(1): p. 57–63.

20. McIlwain, D.R., T. Berger, and T.W. Mak, Caspase functions in cell death and disease. Cold Spring Harb Perspect Biol, 2013. 5(4): p. a008656.

21. Zhirnov, O.P. and V.V. Syrtzev, Influenza virus pathogenicity is determined by caspase cleavage motifs located in the viral proteins. J Mol Genet Med, 2009. 3(1): p. 124–32.

22. Martin, S.J., Caspases: Executioners of Apoptosis, in Pathobiology of Human Disease, L.M. McManus and R.N. Mitchell, Editors. 2014, Academic Press: San Diego. p. 145-152.

23. Nguyen, P.A., et al., pH-induced conformational change of the influenza M2 protein C-terminal domain. Biochemistry, 2008. 47(38): p. 9934–6.

24. Wohlgemuth, N., A.P. Lane, and A. Pekosz, Influenza A Virus M2 Protein Apical Targeting Is Required for Efficient Virus Replication. J Virol, 2018. 92(22).

25. Sanchez-Garrido, J., V. Sancho-Shimizu, and A.R. Shenoy, Regulated proteolysis of p62/SQSTM1 enables differential control of autophagy and nutrient sensing. Sci Signal, 2018. 11(559).

26. Timmer, J.C., et al., Structural and kinetic determinants of protease substrates. Nat Struct Mol Biol, 2009. 16(10): p. 1101–8.

27. Mahrus, S., et al., Global sequencing of proteolytic cleavage sites in apoptosis by specific labeling of protein N termini. Cell, 2008. 134(5): p. 866–76.

28. Dagbay, K.B. and J.A. Hardy, Multiple proteolytic events in caspase-6 self-activation impact conformations of discrete structural regions. Proceedings of the National Academy of Sciences, 2017. 114(38): p. E7977–E7986.

29. Slee, E.A., C. Adrain, and S.J. Martin, Executioner Caspase-3, -6, and -7 Perform Distinct, Non-redundant Roles during the Demolition Phase of Apoptosis*. Journal of Biological Chemistry, 2001. 276(10): p. 7320–7326.

30. Nayak, D.P., et al., Influenza virus morphogenesis and budding. Virus Res, 2009. 143(2): p. 147–61.

31. Iwatsuki-Horimoto, K., et al., The cytoplasmic tail of the influenza A virus M2 protein plays a role in viral assembly. J Virol, 2006. 80(11): p. 5233–40.

32. Schmidt, N.W., et al., Influenza virus A M2 protein generates negative Gaussian membrane curvature necessary for budding and scission. J Am Chem Soc, 2013. 135(37): p. 13710–9.

33. Pena-Francesch, M., et al., The autophagy machinery interacts with EBV capsids during viral envelope release. Proc Natl Acad Sci U S A, 2023. 120(34): p. e2211281120.

34. Arnoldi, F., et al., Rotavirus increases levels of lipidated LC3 supporting accumulation of infectious progeny virus without inducing autophagosome formation. PLoS One, 2014. 9(4): p. e95197.

35. Taisne, C., et al., Human cytomegalovirus hijacks the autophagic machinery and LC3 homologs in order to optimize cytoplasmic envelopment of mature infectious particles. Scientific Reports, 2019. 9(1): p. 4560.

36. Münz, C., The Autophagic Machinery in Viral Exocytosis. Front Microbiol, 2017. 8: p. 269.

37. Nowag, H., et al., Macroautophagy Proteins Assist Epstein Barr Virus Production and Get Incorporated Into the Virus Particles. EBioMedicine, 2014. 1(2-3): p. 116–25.

38. Badham, M.D. and J.S. Rossman, Filamentous Influenza Viruses. Curr Clin Microbiol Rep, 2016. 3(3): p. 155–161.

39. Chu, C.M., I.M. Dawson, and W.J. Elford, FILAMENTOUS FORMS ASSOCIATED WITH NEWLY ISOLATED INFLUENZA VIRUS. The Lancet, 1949. 253(6554): p. 602–603.

40. Wit, E.d., et al., Efficient generation and growth of influenza virus A/PR/8/34 from eight cDNA fragments. Virus Research, 2004. 103(1): p. 155–161.

41. Rogov, V.V., et al., A universal expression tag for structural and functional studies of proteins. Chembiochem, 2012. 13(7): p. 959–63.

42. Kabsch, W., XDS. Acta Crystallogr D Biol Crystallogr, 2010. 66(Pt 2): p. 125–32.

43. Emsley, P., et al., Features and development of Coot. Acta Crystallogr D Biol Crystallogr, 2010. 66(Pt 4): p. 486–501.

44. Winn, M.D., et al., Overview of the CCP4 suite and current developments. Acta Crystallogr D Biol Crystallogr, 2011. 67(Pt 4): p. 235–42.

45. Adams, P.D., et al., PHENIX: a comprehensive Python-based system for macromolecular structure solution. Acta Crystallogr D Biol Crystallogr, 2010. 66(Pt 2): p. 213–21.

46. Hutchinson, E.C. and M. Stegmann, Purification and Proteomics of Influenza Virions. Methods Mol Biol, 2018. 1836: p. 89–120.

47. Demichev, V., et al., DIA-NN: neural networks and interference correction enable deep proteome coverage in high throughput. Nature Methods, 2020. 17(1): p. 41–44.

48. Wallace, A.C., R.A. Laskowski, and J.M. Thornton, LIGPLOT: a program to generate schematic diagrams of protein-ligand interactions. Protein Eng, 1995. 8(2): p. 127–34.

49. Bao, Y., et al., The influenza virus resource at the National Center for Biotechnology Information. J Virol, 2008. 82(2): p. 596–601.

50. Crooks, G.E., et al., WebLogo: a sequence logo generator. Genome Res, 2004. 14(6): p. 1188–90.

51. Schickli, J.H., et al., Plasmid-only rescue of influenza A virus vaccine candidates. Philos Trans R Soc Lond B Biol Sci, 2001. 356(1416): p. 1965–73.

